# 1-[(4-Nitrophenyl)sulfonyl]-4-phenylpiperazine Treatment After Brain Irradiation Preserves Cognitive Function in Mice

**DOI:** 10.1101/2020.01.04.894865

**Authors:** Kruttika Bhat, Paul Medina, Ling He, Le Zhang, Mohammad Saki, Angeliki Ioannidis, Nhan T. Nguyen, Sirajbir S. Sodhi, David Sung, Clara E. Magyar, Linda M. Liau, Harley I. Kornblum, Frank Pajonk

## Abstract

**Background:** Normal tissue toxicity to the CNS is an inevitable consequence of a successful radiotherapy of brain tumors or cancer metastases to the CNS. Cranial irradiation commonly leads to neurocognitive deficits that manifest months or years after treatment. Mechanistically, radiation-induced loss of neural stem/progenitor cells, neuro-inflammation and de-myelinization are contributing factors that lead to progressive cognitive decline.

**Methods:** The effects of Compound #5 on irradiated murine neurospheres, microglia cells and patients-derived gliomaspheres were assessed in sphere-formation assays, flow cytometry and IL-6 ELISAs, Activation of the Hedgehog pathway was studied by qRT-PCR. The *in vivo* effects of Compound #5 were analyzed using flow cytometry, sphere-formation assays, immune-histochemistry, behavioral testing and an intracranial mouse model of glioblastoma.

**Results:** We report that 1-[(4-Nitrophenyl)sulfonyl]-4-phenylpiperazine (Compound #5) mitigates radiation-induced normal tissue toxicity in the brains of mice. Compound #5 treatment significantly increased the number of neural stem/progenitor cells after brain irradiation in female animals, inhibited radiation-induced microglia activation and expression of the pro-inflammatory cytokine interleukin-6. Behavioral testing revealed that treatment with Compound #5 after radiotherapy successfully mitigates radiation-induced decline in motor, sensory and memory function of the brain. In mouse models of glioblastoma, Compound #5 showed no toxicity and did not interfere with the growth-delaying effects of radiation.

**Conclusions:** We conclude that Compound #5 has the potential to mitigate cognitive decline in patients undergoing partial or whole brain irradiation without promoting tumor growth and that the use of this compound as a radiation mitigator of radiation late effects on the CNS warrants further investigation.

**Importance of the Study:** Successful radiotherapy of CNS malignancies inevitably lead to cognitive decline in cancer survivors and treatment options to mitigate this side effect are limited. We present evidence that a piperazine compound can prevent cognitive decline in mice after total brain irradiation without compromising the antitumor effect of radiation, suggesting that this compound could be used to mitigate radiation side effects in brain tumor patients undergoing radiotherapy.

## Introduction

Exposure of the central nervous system to ionizing radiation results in normal tissue toxicity ^1^. With survival times for cancer patients steadily increasing over the past decades ^2^, more and more patients are now at risk of experiencing late effects of radiotherapy. Patients receiving cranial irradiation –and among those in particular pediatric patients– are facing a decline in cognitive function later in life^3-5^. The underlying mechanisms include neuroinflammation, diminished neuronal connectivity and demyelination ^1^. Earlier studies by Limoli and colleagues demonstrated that the functional consequences of brain irradiation can be mitigated by injection of neural stem cells into the brain and that newly derived neurons from this stem cell population integrate into the circuitry of the adult brain ^6^. Furthermore, that activation of microglia is a critical factor for radiation-induced neuroinflammation, which ultimately leads to cognitive decline. These data indicated that mitigating radiation effects in the intrinsic neural stem/progenitor cell population and microglia cells could be exploited in the radiotherapy setting to prevent radiation-induced cognitive decline.

We previously reported that 1-[(4-Nitrophenyl)sulfonyl]-4-phenylpiperazine (compound #5) prevents the acute radiation syndrome in mice by activating the hedgehog signaling pathway ^7, 8^. In this study we tested if compound #5 has an effect on the neural stem/progenitor cell population. Our data show that compound #5 when given after cranial irradiation preserves the neural stem/progenitor cell population, inhibits microglia activation, mitigates radiation-induced neuroinflammation and prevents radiation-induced cognitive impairment in mice.

### Materials and Methods

#### Animals

Nestin-EGFP mice were a kind gift from Dr. Grigori Enikolopov, Cold Spring Harbor Laboratory ^9^. C3Hf/Sed/Kam were originally obtained from the MD Anderson Cancer Center. All experiments were performed in accordance to all local and national guidelines for the care of animals.

For orthotopic tumor grafting 2×10^5^ GL261-Luciferase cells were implanted into the right striatum of the brains of female C57Bl/6 mice using a stereotactic frame (Kopf Instruments, Tujunga, CA) and a nano-injector pump (Stoelting, Wood Dale, IL). Injection coordinates were 0.5 mm anterior and 2.25 mm lateral to the bregma, at a depth of 3.5 mm from the surface of the brain. Tumors were grown for 7 days after which successful grafting was confirmed by bioluminescence imaging.

#### Cell culture

For neural stem cell cultures the entire brains of the newborn Nestin-GFP mice were harvested, minced using a scalpel and transferred to a fresh Eppendorf tube with 1 ml of 0.05 % trypsin-EDTA (Thermo Fisher Scientific, # 25300-054), mixed and incubated at 37°C for 7 minutes. The trypsin was inhibited by adding 1 ml of trypsin inhibitor (Sigma-Aldrich, Cat# T6522). The tissue was centrifuged at 110 x g for 5 minutes and the supernatant was discarded. The tissue was resuspended in 200 µl of NeuroCult (Stem Cell Technologies, # 05702). The solution was mixed well to break down the tissue, diluted with 10 ml NeuroCult media and passed through a 70 µm cell strainer. The cell suspension was centrifuged at 110 x g for 5 minutes, the supernatant discarded and the cell pellet resuspended in 2 ml of ACK lysing buffer (Lonza, Cat# 10-548E). After centrifugation, cells from the final pellet were cultured in 1X complete NeuroCult media (45 ml of base NeuroCult media, 5 ml of supplement media, 1 µl of EGF, 20 µl of FGF, 3130 U/ml of heparin) under standard conditions (37° C, 5% CO_2_) and labeled as Passage #1. Passage #2 cells were used for all experiments.

For microglia cell cultures EOC 20 cells (female; sourced from C3H/HeJ mice) were obtained from ATCC (Manassas, VA). Cells were maintained under standard conditions in DMEM media/10%FBS supplemented with 20% conditioned media from CSF-1-expressing LADMAC bone marrow cells (ATCC).

The primary human glioma cell lines were established at UCLA as described (Hemmati et al., PNAS 2003 ^10^; Characteristics of specific gliomasphere lines can be found in Laks et al., Neuro-Oncology 2016 ^11^). The GL261 murine glioma cell line was from obtained from Charles River

Laboratories, Inc., Frederick, MD. GL261 cells were cultured in log-growth phase in DMEM (Invitrogen, Carlsbad, CA) supplemented with 10% fetal bovine serum, penicillin and streptomycin. Primary HK-374 HK-157, HK-382 glioblastoma cells were propagated as gliomaspheres under serum-free conditions in ultra-low adhesion plates in DMEM/F12, supplemented with B27, EGF, bFGF and heparin as described previously ^10-12^. All cells were grown in a humidified atmosphere at 37°C with 5% CO_2_. The unique identity of all patient-derived specimens was confirmed by DNA fingerprinting (Laragen, Culver City, CA). All lines were routinely tested for mycoplasma infection (MycoAlert, Lonza).

#### Quantitative Reverse Transcription-PCR

Total RNA was isolated using TRIZOL Reagent (Invitrogen). cDNA synthesis was carried out using the SuperScript Reverse Transcription IV (Invitrogen). Quantitative PCR was performed in the QuantStudio 3 (Applied Biosystems, Thermo Fisher, Waltham, MA) using the PowerUp™ SYBR™ Green Master mix (Applied Biosystems, # A25742). *C*_t_ for each gene was determined after normalization to HPRT (mouse) and GAPDH (human) and ΔΔ*C*_t_ was calculated relative to the designated reference sample. Gene expression values were then Ct set equal to 2^−ΔΔC_t_^ as described by the manufacturer of the kit (Applied Biosystems). All PCR primers were synthesized by Invitrogen and designed for the murine and human sequences of Ptch1, Ptch2, Gli1, Gli2, and the housekeeping genes HPRT and GAPDH (**Supplement 1**).

#### Il-6 Enzyme-Linked Immunosorbent Assays

Enzyme-linked immunosorbent assays (ELISA) were performed by following the manufacturer’s instructions (mouse Il-6 Quantikine ELISA Kit, Fisher Scientific, # M6000B.). The absorbance was read at 450 nm (Spectramax M5, Molecular Devices, Sunnyvale, CA). A wavelength correction was performed by subtracting readings at 600 nm from the readings at 450 nm.

#### Irradiation

Neurosphere cultures were irradiated with 0, 2 or 4 Gy at room temperature using an experimental X-ray irradiator (Gulmay Medical Inc. Atlanta, GA) at a dose rate of 5.519 Gy/min. Control samples were sham-irradiated. The X-ray beam was operated at 300 kV and hardened using a 4 mm Be, a 3 mm Al, and a 1.5 mm Cu filter.

Eight-week-old mice were anaesthetized with isoflurane, cone beam computed tomography images were acquired, and individual treatment plans were calculated for each mouse using the SmART Plan software package. Subsequently, the right hemisphere of the brain was irradiated with 4 Gy or 10 Gy using a single beam (for the *in vivo* experiments). For behavioral studies, the whole brain was irradiated with 10Gy using two opposing beams. The X-ray beam was operated at 225 kV. NIST-traceable dosimetry on both X-ray machines was performed using a micro-ionization chamber.

#### In vitro drug treatment

1-[(4-Nitrophenyl)sulfonyl]-4-phenylpiperazine (Compound #5, Vitascreen, Champaign, IL) was solubilized in DMSO. Three hours after irradiation neurosphere cultures were treated with compound #5 (10µM) or DMSO.

#### In vivo drug administration

*In vivo* neural stem/progenitor experiments: Starting twenty-four hours after irradiation mice received 5 daily sub-cutaneous injections of 5 mg/kg of Compound #5 (Vitascreen, Champaign, IL) dissolved in DMSO/Cremophor EL (Sigma-Aldrich, St. Louis, MO) or DMSO/Cremophor EL only.

Behavioral studies: Starting immediately after 10 Gy whole brain irradiation, mice received 5 daily sub-cutaneous injections of Compound #5 or DMSO/Cremophor EL.

#### In vitro assays with patient-derived glioblastoma specimens

For the assessment of self-renewal *in vitro*, HK-374, HK-157 and HK-382 cells were irradiated with 0, or 4 Gy. Three hours after irradiation, cells were treated with either DMSO or Compound #5. The media was supplemented with DMSO or Compound #5 every other day for two weeks. The number of spheres formed in each treatment groups was normalized against the non-irradiated control.

#### Brain dissociation

Five days after drug treatment, the brains of the mice were harvested and placed on the Acrylic Mouse Brain Slicer Matrix (Zivic Instruments, Cat# BSMAA001-1). Coronal sections from 2 mm anterior to 2 mm posterior of the bregma were cut and the left hemisphere was separated from the right. The brain tissue was minced using a scalpel into very small pieces and the cells were isolated as mentioned in the above method section (Neural stem cell culture). The cells were then used for flow cytometric analysis to assess the percentage of Nestin-GFP^+^ cells in different treatment groups or used to quantify self-renewal capacity in neurosphere formation assays.

#### Behavioral Testing

All of the behavioral experiments (Novel Obeject Recognition (NOR), Object in place (OIP), Fear Conditioning (FC)) were conducted in the Behavioral Testing Core (BTC) at UCLA. A detailed description is provided in **Supplement 1**.

#### Flow cytometry

Passage #2 neurospheres established from the brains of Nestin-GFP mice were harvested, dissociated into single cell suspension as described above. Single cell suspensions were either subjected to FACS (Flow Cytometry Core, Terasaki, BD FACS ARIA) for GFP-high, -medium, and –low cell populations or analyzed for GFP expression using a MACSQuant Analyzer (Miltenyi Biosciences, Auburn, CA) and the FlowJo software package (v10, FlowJo, Ashland, OR).

#### Neurosphere-formation assay

In order to assess self-renewal capacity, passage #2 neurospheres from Nestin-GFP mice were trypsinized and plated in 1X complete NeuroCult media into 96-well non-treated plates, ranging from 1 to 1000 cells/well. Growth factors, EGF and bFGF, were added every 3 days, and the cells were allowed to form neurospheres for 14 days. The number of spheres formed per well was then counted and expressed as a percentage of the initial number of cells plated.

#### Immunohistochemistry

Brains were fixed in formalin, embedded in paraffin, and 4 µm coronal sections were stained for Ki67, GFAP and IBA-1 (Translational Pathology Core Laboratory at UCLA). Briefly, slides were placed in xylenes to remove paraffin, then a series of ethanol, washed in tap water, and incubated in 3% hydrogen peroxide/methanol for 10 minutes. After a wash in distilled water, the slides were incubated for 25 minutes in citrate buffer, pH 6 at 95°C. The slides rinsed in PBST (Phosphate Buffered Saline containing 0.05% Tween-20), and then incubated with a rabbit anti-mouse Ki67 antibody (Cell Signaling, Danvers, MA, #12202, 1:200) or rabbit anti-GFAP antibody (Cell Signaling, #12389S, 1:50) at RT for 1 hr and with a rabbit anti-IBA-1 antibody (Abcam, Cambridge, MA, #ab178847, 1:8000) overnight at 4 °C. The slides were rinsed with PBST, and were incubated with secondary antibody solution (Cell Signaling SignalStain Boost IHC Detection Reagent, HRP, Rabbit) at RT for 30 minutes. Slides were rinsed with PBST and incubated with DAB (3,3’-Diaminobenzidine, Cell Signaling SignalStain DAB Substrate Kit, #8059), washed in tap water, counterstained with Harris’ Hematoxylin, dehydrated in ethanol, and mounted with media.

Slides were digitized on a ScanScope AT (Aperio Technologies, Inc., Vista, CA) and morphimetric analysis performed with *Definiens’* Tissue Studio (Definiens Inc., Parsippany, NJ) for unbiased determination of the percentage of Ki67-, GFAP-, or IBA-1-positive cells. Scanning and analyses were performed through the Translational Pathology Core Laboratory, Department of Pathology and Laboratory Medicine, David Geffin School of Medicine at UCLA.

#### Statistics

All statistical analyses were performed using the Graphpad Prism software package. Unless stated otherwise, results were derived from at least three animals per group. A *p*-value equal to or smaller than 0.05 in a student t-test or one-way ANOVA test was considered statistically significant. Kaplan-Meier estimates were calculated using the GraphPad Prism Software package and a *p*-value of 0.05 in a log-rank test indicated a statistically significant difference.

### Results

#### Radiation mitigation in neural stem/progenitor cells in vitro

Passage #2 neurospheres from Nestin-GFP mice, in which most cells were Nestin-GFP^+^, were used for all *in vitro* experiments (**Figure 1A**). In order to test the self-renewing capacity of the sorted GFP-high, -medium, and –low cells from neurospheres, we performed *in vitro* limiting dilution assays. Nestin-GFP^high^ cells showed 4.3-fold higher sphere-formation than Nestin-GFP^med^ and 13.5-fold higher sphere-formation than Nestin-GFP^low^ cells (**Figure 1B**), thus supporting the neural stem/progenitor phenotype of Nestin-GFP^high^ cells. Irradiation of the neurospheres with 0, 2 or 4 Gy preferentially reduced the size of the Nestin-GFP^high^ cell population (**Figure 1C**). This was in line with a previous report on radiation-induced differentiation of neural stem/progenitor cells ^13^.

**Figure 1.**
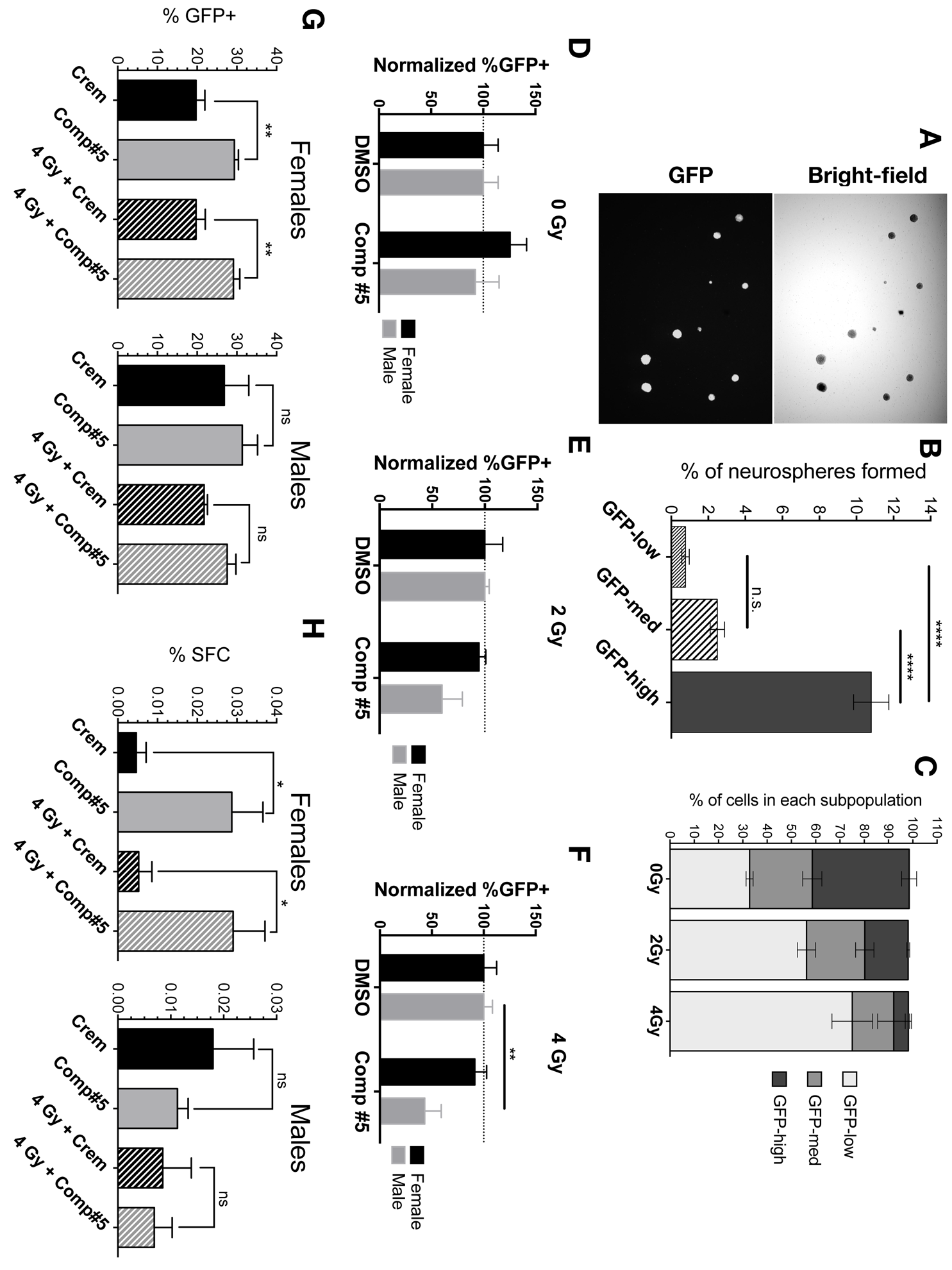
Radiation mitigation in neural/progenitor cells in vitro and in vivo. **(A)** Representative images of neurosphere cultures established from the brains of Nestin-GFP mice. Bright-field and GFP image (4X). **(B)** Sorted GFP^high^, -^medium^ and –^low^ cells were subjected to an *in vitro* limiting dilution assay. **(C)** Effect of radiation (0, 2 or 4 Gy) on three different subpopulations of Nestin-GFP neurospheres. Neurospheres (passage #2) from male or female newborn pups were subjected to 0 (**D**), 2 (**E**) or 4 Gy (**F**) followed by a single treatment with either DMSO or Compound #5 (10 µM) three hours post irradiation. Eight-week old Nestin-GFP male and female mice were sham irradiated or irradiated with 4 Gy. After Twenty-four mice were treated with Cremophor or Compound #5 (5 mg/kg) sub-cutaneously for five days. The brains of the mice were harvested, dissociated and analyzed by FACS (**G**) or sphere forming assays (**H**). All experiments in this figure have been performed with at least 3 independent biological repeats. (Unpaired t-test. * *p*-value < 0.05, ** *p*-value < 0.01, **** *p*-value < 0.0001).

Next, we irradiated neurospheres with 0, 2, or 4 Gy and treated the cells with Compound #5. Compound #5 failed to increase the size of the population of Nestin-GFP^high^ neural stem/progenitor cells (**Figure 1D-F**). While Compound #5 did not show any effect on the neurospheres derived from the female newborn pups, it significantly reduced the Nestin-GFP^high^ population of cells derived from the male newborn pups, especially in the 4 Gy treated groups.

#### Radiation mitigation in neural stem and progenitor cells in vivo

We next considered the possibility that our lead compound targets neural stem/progenitor cells indirectly, which cannot be easily tested in the absence of the correct microenvironment *in vitro*. To test this, 8 week-old male and female Nestin-GFP mice were irradiated with a dose of 4 Gy to the right brain hemisphere. The radiation treatment plans ensured sparing of the contralateral hemisphere from irradiation, thus allowing for an internal unirradiated control for each individual mouse. Twenty-four hours later the animals were treated with 4 daily injections of Cremophor or compound #5. Tthe brains were harvested, digested and analyzed for the number of Nestin-GFP^high^ stem/progenitor cells and the cells tested for their self-renewing capacity.

Compound #5 significantly increased the number of Nestin-GFP^high^ stem/progenitor cells in female mice (**Figure 1G**, **left panel**). In male mice we observed a similar trend but the effect was not significant (**Figure 1H, left panel**). In *in vitro* limiting dilution assays we observed a significant increase in sphere-forming capacity in cells obtained from female mice treated with compound #5 but not in cells obtained from male mice (**Figure 1G** & H, **right panel**). Therefore, all remaining studies were conducted in female mice.

#### Compound #5 mitigates radiation-induced neuro-inflammation

Six-week old female C3H mice treated with a single fraction of 4 or 10 Gy to the right brain hemisphere (**Figure 2A**). Starting wenty-four hours after irradiation, the mice were treated with either Cremophor or Compound #5 for 5 days. The brains were harvested, fixed in formalin, embedded in paraffin and 4 µM sections subjected to immunohistochemistry (IHC). Sections were stained with H&E, or against GFAP (reactive astrocytes marker), IBA1 (activated microglia marker) and Ki67 (proliferation marker) and subjected to an automated image analysis (**Figure 2B-E**).

**Figure 2.**
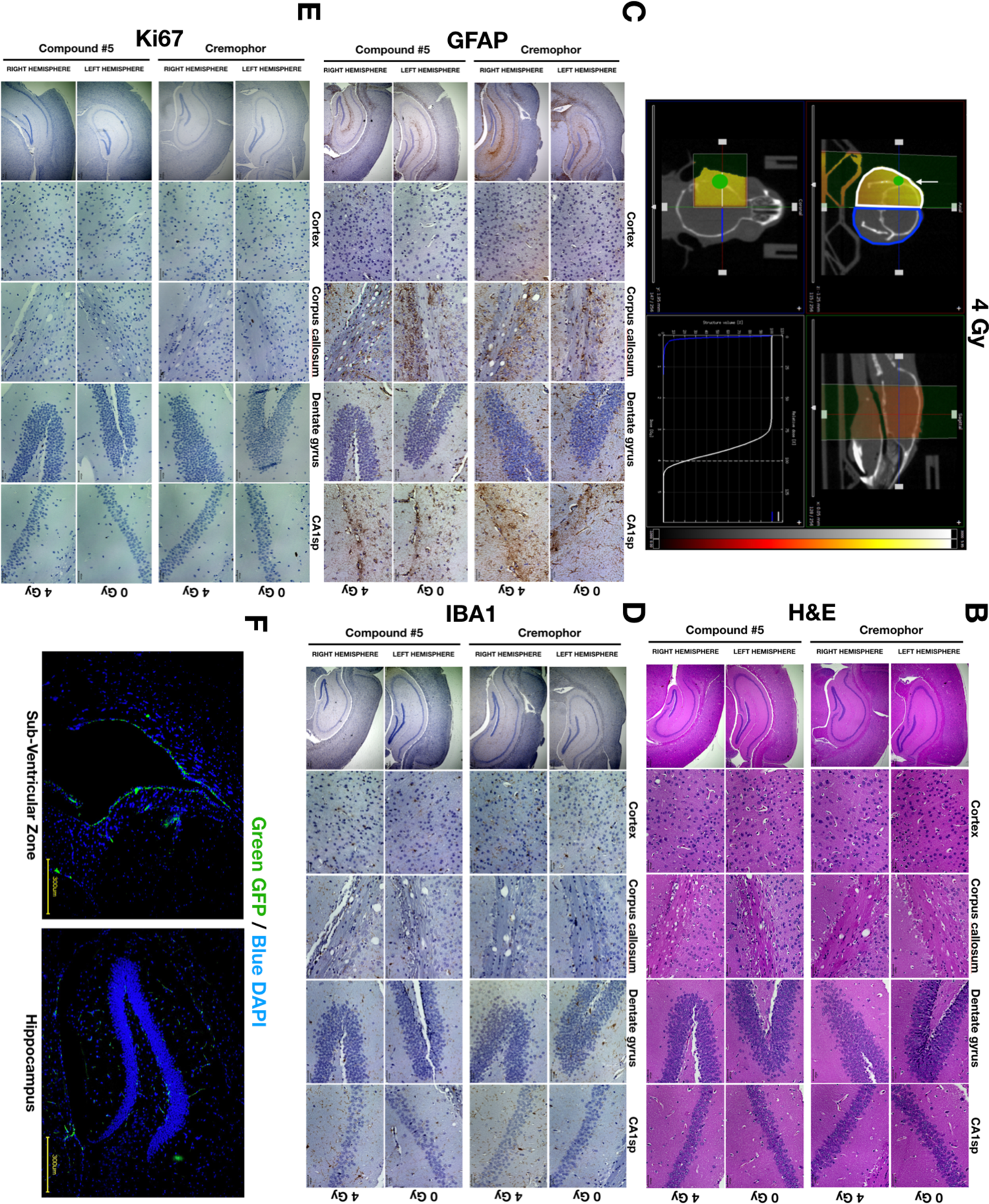
Radiation treatment plans and IHC images. (A) Six-week old C3H female mice were irradiated with 0 or 4 Gy and and treated with Cremophor or Compound #5 (5 mg/kg, s.c.) for five days. Coronal sections were stained with H&E (**B**) or labeled with antibodies against GFAP (**C**), IBA1 (**D**) or Ki67 (**E**). Representative images (10X) of the sub-ventricular zone and hippocampus region of eight-week old Nestin-GFP mice labeled for GFP (**F**).

The slides were scored for positively stained cells in the cortex, corpus callosum, CA1sp, and the dentate gyrus. In the non-irradiated group, Compound #5 did not show significant changes in GFAP, IBA-1, or Ki67 expression. When Compound #5 was combined with radiation it led to a significant reduction in GFAP and IBA1 expression, thus indicating mitigation of radiation-induced neuro-inflammation (**Figure 3A**).

**Figure 3.**
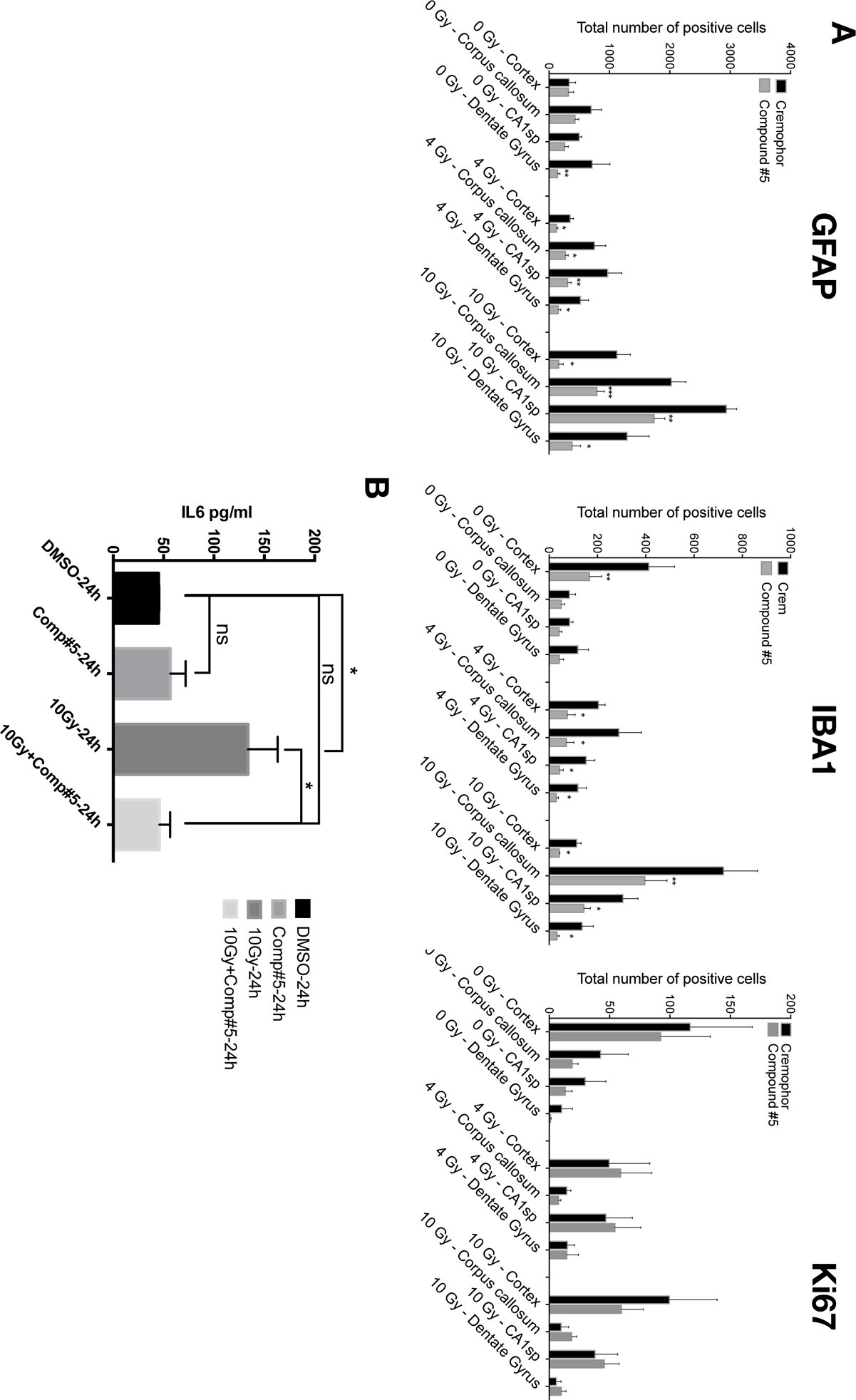
Compound #5 mitigated radiation-induced neuro-inflammation. (A) Coronal sections stained against GFAP (left panel), IBA1 (center panel) and Ki67 (Right panel) were quantified for the total number of positive cells in the cortex, corpus callosum, CA1sp and dentate gyrus regions of the brain in the unirradiated and irradiated mice. (B) ELISA for IL-6 using conditioned media from normal EOC20 microglia cells twenty-four hours after exposing them to 0 or 4 Gy irradiation and treatment with either DMSO or Compound #5. (Unpaired t-test. * *p*-value < 0.05, ** *p*-value < 0.01, *** *p*-value < 0.001).

To further confirm the anti-inflammatory effect of compound #5, we collected conditioned media from EOC20 microglia cells twenty-fours exposure after irradiation with 0 or 10 Gy and treatment with DMSO or compound #5 *in vitro*. IL-6 secretion levels were assessed usiging san ELISA. In line with the well-known pro-inflammatory effect of radiation we observed a significant increase in the secretion of IL-6 in cells treated with 10 Gy. Consistent with an anti-inflammatory effect, Compound #5 significantly reduced IL-6 secretion levels (4 Gy: 2.5-fold, p=0.0297; 10 Gy: 2-fold, p=0.0425) when compared to the corresponding control group (**Figure 3B**, left).

#### Preservation of cognitive function in irradiated mice

Ionizing radiation to the brain has long been known to cause neuroinflammation, which ultimately leads to a decline in cognitive function ^14,15^. Our short-term experiments indicted that Compound #5 mitigated neuroinflammation. Next we sought to test if treatment with compound #5 also translated into improved cognitive function. After total brain irradiation with 10 Gy (**Figure 4A**) animals were treated with either Cremophor or Compound #5 (5 mg/kg) for 5 days. One month after irradiation the animals were subjected to unbiased cognitive testing (**Figure 4B**). Novel Object Recognition (NOR) and Object In Place (OIP) was performed to evaluate impairments in the prefrontal and perirhinal cortices and hippocampus regions. This was followed by Fear Conditioning (FC) tasks for studying deficits in motor, sensory and memory function dependent on the hippocampal regions.

**Figure 4.**
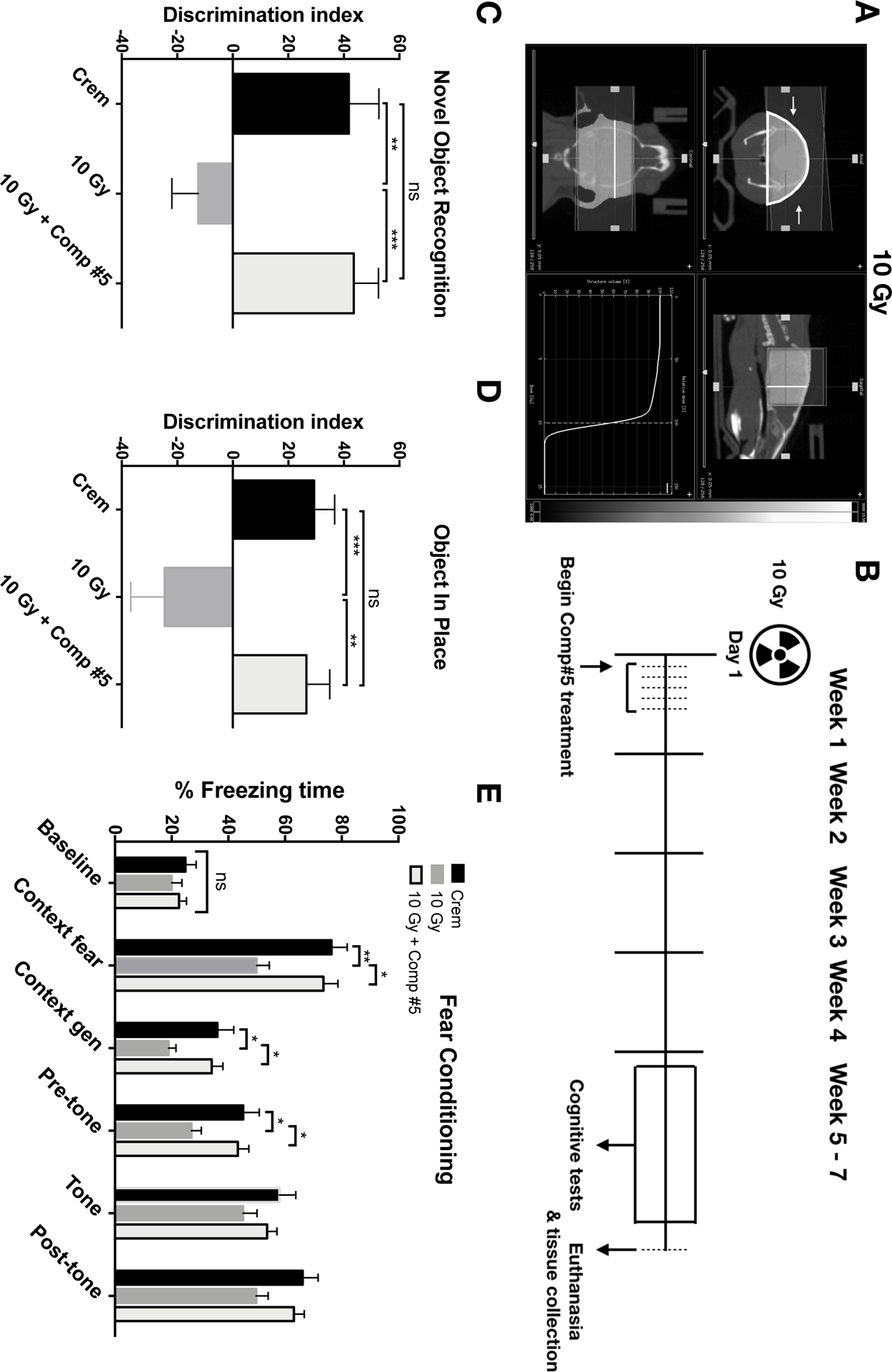
Preservation of cognitive function in irradiated mice. Groups (n=10) of six-week old female C3H mice were irradiated with 0 or 10 Gy (**A**) and treated with Compound #5 (5 mg/kg) or Cremophor for 5 days. Schematic representation of the behavioral testing schedule (**B**). Starting week 5 the mice were tested for cognitive functions by performing Novel Object Recognition (NOR, **C**), Object In Place (OIP, **D**) and Fear Conditioning (FC, **E**) tasks. (Unpaired t-test. * *p*-value < 0.05, ** *p*-value < 0.01, *** *p*-value < 0.001).

Cranial irradiation of 10 Gy demonstrated a significant behavioral deficit on both NOR and OIP tasks compared to control (Cremophor) groups as indicated by impaired preference to novel object (**Figure 4C**) or place (**Figure 4D**); however when combined with Compound #5 the mice showed significantly improved performance in identifying the novel object (**Figure 4C**) or place (**Figure 4D**). Furthermore, the discrimination index (DI) between the un-irradiated and the combined treatment groups were statistically insignificant, indicating that Compound #5 had successfully mitigated the radiation effects. In the FC task, the baseline freezing levels were comparable among the three treatment groups. All groups also showed an increased freezing behavior post 3 tone-shock pairings (context fear bars). Baseline freezing levels forty-eight hours post-trainings showed significantly decreased freezing in irradiated mice when compared to the un-irradiated control mice. Administration of Compound #5 to the irradiated mice reduced the cognitive deficits (**Figure 4E**). Treatment of the irradiated mice with Compound #5 led to an increased freezing behavior when compared to the irradiated group, indicating preservation of hippocampal function (**Figure 4E**).

#### Effects of Compound #5 on GBM cells in vitro and in vivo

Radiation mitigators or protectors always bear the risk of radiation protection or mitigation not only in normal tissues but also in tumors. To test the effect of Compound #5 on glioblastoma cells *in vitro* we performed sphere-forming capacity assays using three different patient-derived GBM cells lines, HK-374, HK-157 and HK-382 in the presence (10 µM) or absence of Compound #5 in combination with irradiation at 0 or 4 Gy. The gliomaspheres were treated with DMSO or Compound #5 every other day for 2 weeks at the end of which the number of spheres formed was counted and presented as percentage of spheres formed. In HK-374 GBM cells Compound #5 significantly reduced its self-renewing capacity with or without irradiation (**Figure 5A**, left panel) while in the HK-157 and HK-382 cell lines Compound #5 had no effect (**Figure 5A**, center and right panel).

**Figure 5.**
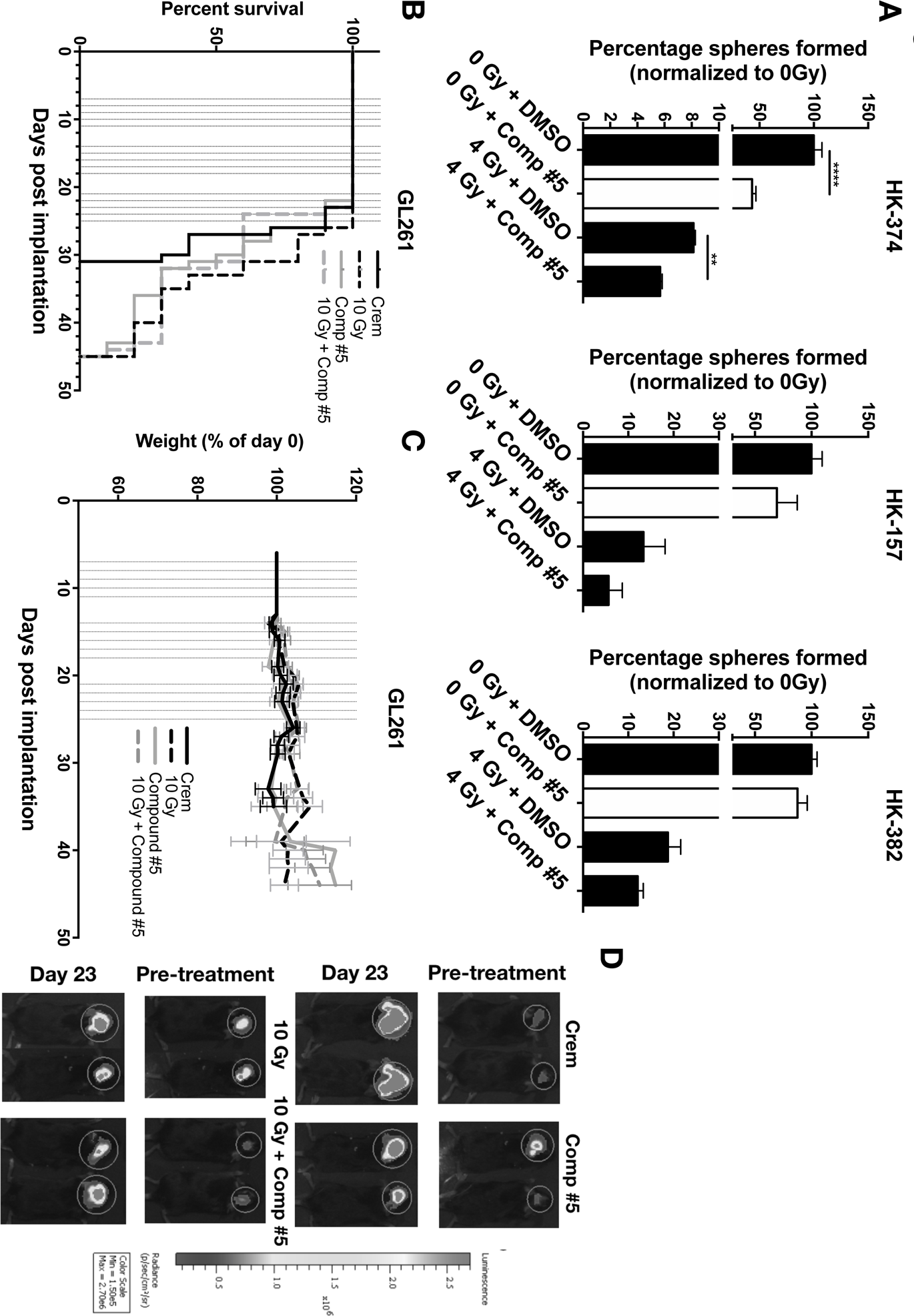
Effects of Compound #5 on GBM cells in vitro and in vivo. **(A)** Patient-derived HK-374 (left panel), HK-157 (center panel) and HK-382 (right panel) GBM cells were used to perform sphere-forming assays with sham-irradiated or irradiated cells in the presence or absence of Compound #5 (10 µM). The cells were treated every other day for two weeks. The number of spheres formed under each condition were counted and presented as percentage spheres formed. (Unpaired t-test. ** p-value < 0.01, **** p-value < 0.0001). (**B**) 2×10^5^ GL261-luciferase mouse glioma cells were implanted intracranially into six-week old female C57BL/6 mice. Animals were irradiated with 0 or 10 Gy and treated with Cremophor or Compound #5 (5 mg/kg, s.c.) for 3 weeks. The effect of Compound #5 on survival in tumor-bearing mice was assessed using Kaplan-Meier estimates. (**C**) Weight curves for the mice in the different treatment groups. (**D**) Bio-luminescence images of mice bearing tumors obtained at Day 7 (pre-treatment) and at Day 23 (days post implantation). Each group had n=10 mice.

To test if Compound #5 interfered with the effects of tumor irradiation *in vivo*, 2×10^5^ GL261-luciferase mouse glioma cells were intracranially injected in C57BL/6 mice. Seven days after implantation, tumor grafting was confirmed by bioluminescence imaging (BLI), and the tumors were either sham irradiated or irradiated with 10 Gy. Immediately after irradiation, the mice were treated with either Cremophor or Compound #5 (5 mg/kg) sub-cutaneously. The treatment was given on a 5-days on / 2 days-off schedule for 3 weeks. Weights of the mice were recorded everyday until the study end point. Kaplan-Meier survival estimates showed no effects of Compound #5 alone or in combination with radiation (**Figure 5B**). Importantly, Compound #5 did not show any toxicity and did not lead to weight loss (**Figure 5C**). Tumor growth was monitored by the bioluminescence imaging of the tumors at Day 7 (pre-treatment) and at Day 23 (days post implantation, **Figure 5D**) and indicated no tumor-promoting effects of Compound #5.

#### Effects of Compound #5 on the Hedgehog pathway in microglia and GBM cells

Recent publications indicated that deregulated developmental pathways play a key role in GBM progression and tumorigenesis by conferring drug resistance to the tumor cells and that inhibition of Hedgehog pathway induced apoptosis in GBM cells ^16-18^. We had previously demonstrated that Compound #5 activates the Hedgehog pathway through binding to the transmembrane domain of Smoothened ^7^. Therefore, we sought to test, if microglia and glioblastoma cells show different sensitivities to Compound #5 that would explain its differential effect in normal tissues and tumors. qRT-PCR for Hedgehog target genes was performed in normal microglia cells (EOC20) and the patient-derived GBM tumor cells (HK-374), 24 hours after treatment with different concentrations of Compound #5. The results obtained are presented as a ratio of fold changes of the genes in EOC20 over HK-374 cells. At low doses of Compound #5 (500 nM to 1 µM) it induced expression of the Hedgehog pathway targets genes Ptch1, Ptch2, Gli1 and Gli2 in microglia cells more efficiently when compared to HK-374 GBM cells, both alone and in combination with radiation (Figure 6A – D), suggesting that microglia cells are more sensitive to Compound #5 than HK-374 glioma cells.

**Figure 6.**
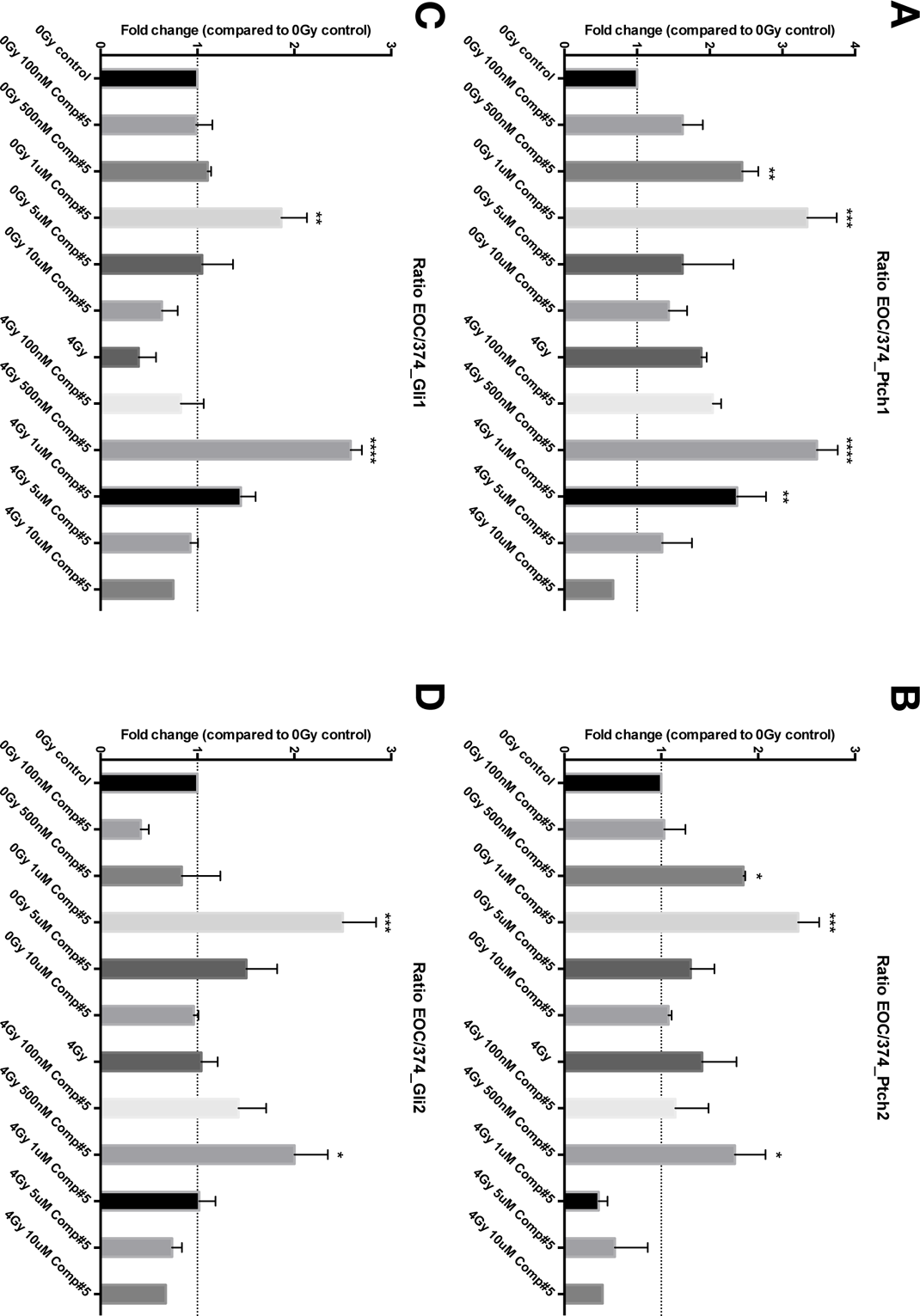
Effects of Compound #5 on Hedgehog signaling in microglia and GBM cells. qRT-PCR for Hedgehog target genes Ptch1 (A), Ptch2 (B), Gli1 (C) and Gli2 (D) in EOC20 microglia and HK-374 glioma cells twenty-fours hours after irradiation and treatment with DMSO or Compound #5. Fold-changes are presented as a ratio of EOC20 microglia cells over HK-374 cells. (Un-paired One-way ANOVA test. * *p*-value < 0.05, ** *p*-value < 0.01, *** *p*-value < 0.001, **** *p*-value < 0.0001).

### Discussion

Aside from surgery, radiotherapy is one of the most effective cancer treatments for patients suffering from brain cancer or cancer metastases to brain. However, with 5-year survival rates steadily increasing, more and more patients experience long-term treatment side effects, which in the case of cranial irradiation manifest in impaired cognitive function. Symptoms arise months and years after completion of radiotherapy and are particularly detrimental in childhood cancer survivors where tumor control rates are often excellent and the cognitive decline can amount to a loss of 1-2 IQ points per year ^19^.

Some experimental approaches, while difficult to translate into the clinic, have shown promising results ^20-22^, but approved clinical treatment options for preventing the late sequelae of cerebral radiotherapy are few and are mostly limited to radiation treatment volume reduction ^23^ or sparing of critical brain structures from irradiation ^24^. Previous pharmacological radioprotection studies using e.g. Amifostine have been hindered by the lack blood-brain-barrier penetration of the drugs and the general concern of simultaneous protection of tumor cells ^25^.

Few pharmacological treatment attempts have been made to mitigate radiation effects to CNS the after completion of treatment and were mostly limited to corticosteroids, which are routinely used to acutely reduce edema but are not sustainable as a long-term treatment option. So far experimental approaches had limited ^26^ or no success ^27^.

We had previously reported that 1-[(4-Nitrophenyl)sulfonyl]-4-phenylpiperazine (compound #5) mitigated the acute intestinal radiation syndrome when given 24 hours or later after a lethal dose of radiation through activation of the hedgehog pathway ^7,8^. Motivated by reports in the literature that hedgehog signaling also affects neural stem cells ^28-30^ we sought to test if compound #5 would mitigate radiation injury in brain tissues.

When given after total brain irradiation compound #5 increased the number of Nestin-GFP-positive cells and their self-renewal capacity in the brains of female mice while it had no effect on male mice. It is noteworthy that the self-renewal of Nestin-GFP-positive cells from male mice exceeded that of female mice, both at baseline and after 4 Gy and that the number and self-renewal capacity of Nestin-GFP-positive cells in female mice were not affected by a single dose of 4 Gy. Our data does not explain the striking gender differences in efficacy for Compound #5 but it underlines the need of testing of compounds in both genders.

Attempts to show the effect of compound #5 on passage #2 neural stem/progenitor cells *in vitro* failed irrespective of gender, indicating that compound #5 does not have a direct effect on neural stem/progenitor cells but that it rather affects the microenvironment. The possibility of indirect effects was further supported by data showing a reduction of radiation-induced IL-6 production by microglia cells *in vitro* and reduction of radiation-induced astrogliosis (GFAP) and microglia activation (IBA1) *in vivo*.

Importantly, compound #5 was well tolerated with no side effects observed and it did not attenuate the effects of radiation on glioma cells *in vitro* or *in vivo*, This indicates that compound #5 could be safely administered during or after completion of radiotherapy in patients suffering from glioblastoma, where the presence of residual tumors cells after completion of surgery and radiotherapy is almost always inevitable. An explanation for the differential effects of compound #5 on normal and malignant cells could be that the Hedgehog signaling pathway is utilized in gliomas at a different threshold resulting in differential responses to activators of this pathway. Lastly, the observed effects of compound #5 were not restricted to changes at the cellular level but translated into preservation of cognitive function in the animals.

We conclude, that 1-[(4-Nitrophenyl)sulfonyl]-4-phenylpiperazine or its analogs have the potential to mitigate radiation effects to the normal brain when given during or after radiotherapy and warrant further investigation.

## Notes

**Funding**: FP, LML and HIK were supported by grants from the *National Cancer Institute* (P50CA211015, R01CA200234) and the National Institute of Allergies and Infectious Diseases (U19AI067769).

**Conflict of Interest:** The authors declare no conflict of interest.

## References

1. Jellinger K. Human central nervous system lesions following radiation therapy. Zentralbl Neurochir. 1977; 38(2):199–200.

2. SEER. Cancer Stat Facts: Brain and Other Nervous System Cancer. https://seer.cancer.gov/statfacts/html/brain.html.

3. Tso WWY, Liu APY, Lee TMC, et al. Neurocognitive function, performance status, and quality of life in pediatric intracranial germ cell tumor survivors. J Neurooncol. 2019; 141(2):393–401.

4. Szentes A, Eros N, Kekecs Z, et al. Cognitive deficits and psychopathological symptoms among children with medulloblastoma. Eur J Cancer Care (Engl). 2018; 27(6):e12912.

5. Doger de SpevilleE, Robert C, Perez-Guevara M, et al. Relationships between Regional Radiation Doses and Cognitive Decline in Children Treated with Cranio-Spinal Irradiation for Posterior Fossa Tumors. Front Oncol. 2017; 7:166.

6. Smith SM, Limoli CL. Stem Cell Therapies for the Resolution of Radiation Injury to the Brain. Curr Stem Cell Rep. 2017; 3(4):342–347.

7. Duhachek-Muggy S, Bhat K, Medina P, et al. Radiation Mitigation of the Intestinal Acute Radiation Injury in Mice by 1-[(4-Nitrophenyl)Sulfonyl]-4-Phenylpiperazine. Stem Cells Transl Med. 2019.

8. Bhat K, Duhachek-Muggy S, Ramanathan R, et al. 1-(4-nitrobenzenesulfonyl)-4-phenylpiperazine increases the number of Peyer’s patch-associated regenerating crypts in the small intestines after radiation injury. Radiother Oncol. 2019; 132:8–15.

9. Mignone JL, Kukekov V, Chiang AS, Steindler D, Enikolopov G. Neural stem and progenitor cells in nestin-GFP transgenic mice. J Comp Neurol. 2004; 469(3):311–324.

10. Hemmati HD, Nakano I, Lazareff JA, et al. Cancerous stem cells can arise from pediatric brain tumors. Proc Natl Acad Sci U S A. 2003; 100(25):15178–15183.

11. Laks DR, Crisman TJ, Shih MY, et al. Large-scale assessment of the gliomasphere model system. Neuro Oncol. 2016; 18(10):1367–1378.

12. Vlashi E, Kim K, Lagadec C, et al. In vivo imaging, tracking, and targeting of cancer stem cells. Journal of the National Cancer Institute. 2009; 101(5):350–359.

13. Konirova J, Cupal L, Jarosova S, et al. Differentiation Induction as a Response to Irradiation in Neural Stem Cells In Vitro. Cancers (Basel). 2019; 11(7).

14. Parihar VK, Acharya MM, Roa DE, Bosch O, Christie LA, Limoli CL. Defining functional changes in the brain caused by targeted stereotaxic radiosurgery. Transl Cancer Res. 2014; 3(2):124–137.

15. Acharya MM, Green KN, Allen BD, et al. Elimination of microglia improves cognitive function following cranial irradiation. Sci Rep. 2016; 6:31545.

16. Wang K, Chen D, Qian Z, Cui D, Gao L, Lou M. Hedgehog/Gli1 signaling pathway regulates MGMT expression and chemoresistance to temozolomide in human glioblastoma. Cancer Cell Int. 2017; 17:117.

17. Melamed JR, Morgan JT, Ioele SA, Gleghorn JP, Sims-Mourtada J, Day ES. Investigating the role of Hedgehog/GLI1 signaling in glioblastoma cell response to temozolomide. Oncotarget. 2018; 9(43):27000–27015.

18. Ulasov IV, Nandi S, Dey M, Sonabend AM, Lesniak MS. Inhibition of Sonic hedgehog and Notch pathways enhances sensitivity of CD133(+) glioma stem cells to temozolomide therapy. Mol Med. 2011; 17(1-2):103–112.

19. Kahalley LS, Ris MD, Grosshans DR, et al. Comparing Intelligence Quotient Change After Treatment With Proton Versus Photon Radiation Therapy for Pediatric Brain Tumors. J Clin Oncol. 2016; 34(10):1043–1049.

20. Leavitt RJ, Limoli CL, Baulch JE. miRNA-based therapeutic potential of stem cell-derived extracellular vesicles: a safe cell-free treatment to ameliorate radiation-induced brain injury. Int J Radiat Biol. 2019; 95(4):427–435.

21. Hinzman CP, Baulch JE, Mehta KY, et al. Plasma-derived extracellular vesicles yield predictive markers of cranial irradiation exposure in mice. Sci Rep. 2019; 9(1):9460.

22. Baulch JE, Acharya MM, Allen BD, et al. Cranial grafting of stem cell-derived microvesicles improves cognition and reduces neuropathology in the irradiated brain. Proc Natl Acad Sci U S A. 2016; 113(17):4836–4841.

23. Brown PD, Jaeckle K, Ballman KV, et al. Effect of Radiosurgery Alone vs Radiosurgery With Whole Brain Radiation Therapy on Cognitive Function in Patients With 1 to 3 Brain Metastases: A Randomized Clinical Trial. JAMA. 2016; 316(4):401–409.

24. Gondi V, Pugh SL, Tome WA, et al. Preservation of memory with conformal avoidance of the hippocampal neural stem-cell compartment during whole-brain radiotherapy for brain metastases (RTOG 0933): a phase II multi-institutional trial. J Clin Oncol. 2014; 32(34):3810–3816.

25. Lindegaard JC, Grau C. Has the outlook improved for amifostine as a clinical radioprotector? Radiother Oncol. 2000; 57(2):113–118.

26. Brown PD, Pugh S, Laack NN, et al. Memantine for the prevention of cognitive dysfunction in patients receiving whole-brain radiotherapy: a randomized, double-blind, placebo-controlled trial. Neuro Oncol. 2013; 15(10):1429–1437.

27. Rapp SR, Case LD, Peiffer A, et al. Donepezil for Irradiated Brain Tumor Survivors: A Phase III Randomized Placebo-Controlled Clinical Trial. J Clin Oncol. 2015; 33(15):1653–1659.

28. Kume M, Chiyoda H, Kontani K, Katada T, Fukuyama M. Hedgehog-related genes regulate reactivation of quiescent neural progenitors in Caenorhabditis elegans. Biochem Biophys Res Commun. 2019; 520(3):532–537.

29. Klein SD, Nguyen DC, Bhakta V, et al. Mutations in the sonic hedgehog pathway cause macrocephaly-associated conditions due to crosstalk to the PI3K/AKT/mTOR pathway. Am J Med Genet A. 2019; 179(12):2517–2531.

30. Yang H, Liu C, Fan H, et al. Sonic Hedgehog Effectively Improves Oct4-Mediated Reprogramming of Astrocytes into Neural Stem Cells. Mol Ther. 2019; 27(8):1467–1482.

