## Supplementary Material for "1-[(4-Nitrophenyl)sulfonyl]-4-phenylpiperazine Treatment After Brain Irradiation Preserves Cognitive Function in Mice"

**Primer sequences**

mouse primers

Ptch1

For: CTTCTCCTATCTTCTGACGGGGT

Rev: AAAGAACTGCGGCAAGTTTTTG

Ptch2

For: TGCAGAGCACCTTCCTG

Rev: CGATGTCATGTGTCTGGTAG

Gli1

For: CCGACGGAGGTCTCTTTGTC

Rev: AACATGGCGTCTCAGGGAAG

Gli2

For: AAGCACCAGAACCGCACTCACTC

Rev: CTTGAGCAGTGGAGCACGGACAT

HPRT

For: GCTGGTGAAAAGGACCTCT

Rev: CACAGGACTAGAACACCTGC

human primers

Ptch1

For: CTGCGTCAGCAGAGTGATTC

Rev: AGCTGAGGGTGTCCTGTGTC

Ptch2

For: GGAATGATTGAGCGGATGATTGA

Rev: CCACCTGTGCCTTGTCTAGC

Gli1

For: CCACGGGGAGCGGAAGGAG

Rev: ACTGGCATTGCTGAAGGCTTTACTG

Gli2

For: TGACATTCGGCTAACGAGGG

Rev: CCCAGACTCGGCTTTACGGA

GAPDH

For: CCCACTCCTCCACCTTTGA

Rev: TGGTGGTCCAGGGGTCTT

**Behavioral Testing**

Mice were transferred to the BTC at seven-weeks and allowed to acclimate for 1 week. The following behavioral tests were then performed.

Fear Conditioning: Mice were handled for 5 days prior to beginning the experiment to acclimate them to the experimenter. On all three days of the experiment, mice were transferred to a holding room and allowed to acclimate for at least 30 minutes. They were transferred to the experiment room in individual clean empty plastic tub cages with a lid on a utility cart. The lid type was changed on Day 3 to create a different environment.

On day 1 of the experiment, mice were placed in the standard configuration of the MedAssociates Fear Conditioning mouse chambers which were cleaned with 70% Isopropyl alcohol to deodorize between trials and scented with 50% Windex. The fan inside the chamber was kept on. After a 180s acclimation period, all the mice received 3 presentations a 30 second 2800 Hz tone co-terminating with a 2 second 0.75mA shock with an inter-shock interval of 180 seconds. They were removed from the chambers 30 seconds after the final shock and returned to their home cages.

On day 2, the context test was performed: mice were returned to the same environment as they were trained with on day 1 and were placed in the chambers for 8 minutes with no stimuli.

On day 3, mice were placed in an altered environment which had plastic flooring covering the shock grid, a plastic curved wall insert to change the shape of the environment, had the fan off, was cleaned with Ethanol and scented with 25% Simple Green. The light was kept off for the same reason as on day 1. The location of each subject was changed from the previous day. Mice received the same procedure as on day 1 except that there were no shocks.

All the data was evaluated by MedAssociates with a motion threshold set to 18. Results displayed are percent time freezing during each component of the experiment.

Novel Object Recognition (NOR) / Object In Place (OIP): The mice were run in the following manner: The first 2 days, mice were exposed to the open arena (30x30 cm square) for 10 min to habituate to the environment. For NOR, mice were exposed to two identical objects for 10 min. They were then removed and placed in a holding cage for 5 min. One object was replaced with a novel object, and the mice were placed back in the chamber for 10 min. This procedure was repeated for the OIP, but instead of replacing an object, one of the identical objects was moved to a new location. Completely different objects were used for the NOR and the OIP.
